# Aquaporins are main contributors of root hydraulic conductivity in pearl millet [*Pennisetum glaucum* (L) R. Br.]

**DOI:** 10.1101/2020.05.13.094094

**Authors:** Alexandre Grondin, Pablo Affortit, Christine Tranchant-Dubreuil, Carla de la Fuente Cantó, Cédric Mariac, Pascal Gantet, Vincent Vadez, Yves Vigouroux, Laurent Laplaze

## Abstract

Pearl millet is a key cereal for food security in arid and semi-arid regions but its yield is increasingly threatened by water stress. Physiological mechanisms consisting in saving water or increasing water use efficiency can alleviate that stress. Aquaporins (AQP) are water channels contributing to plant hydraulic balance that are supposedly involved in these mechanisms by mediating root water transport. However, AQP remain largely uncharacterized in pearl millet. Here, we studied AQP function in root water transport in two pearl millet lines contrasting for water use efficiency (WUE). We observed that these lines were also contrasting for root hydraulic conductivity (Lpr) and AQP contribution to Lpr, the line with lower WUE showing significantly higher AQP contribution to Lpr. To investigate the AQP isoforms contributing to Lpr, we developed genomic approaches to first identify the entire AQP family in pearl millet and second study the plasma membrane intrinsic proteins (PIP) gene expression profile. We identified and annotated 33 AQP genes in pearl millet among which ten encoded PIP isoforms. *PgPIP1-3* and *PgPIP1-4* were significantly more expressed in the line showing lower WUE, higher Lpr and higher AQP contribution to Lpr. Overall, our study suggests that AQP from the PIP1 family are the main contributor of Lpr in pearl millet and are possibly associated to whole plant water use mechanisms. This study paves the way for further investigations on AQP functions in pearl millet hydraulics and adaptation to environmental stresses.

The newly sequenced nucleotide sequences reported in this article have been submitted to GenBank under the submission number 2333840 (TPA grp467567). Assignment of GenBank accession number is in process.

## Introduction

Plant hydraulics depends on soil water capture by roots, transport to the leaves and diffusion as vapor from the stomatal cavity to the atmosphere. In this plant hydraulic continuum, the radial water transport from the soil solution to the xylem vessels uses two paths: the apoplastic path where water can flow along the cell wall structures, and the cell to cell path where water can flow across cell membranes (transcellular) or along cytoplasmic continuities formed by plasmodesmata (symplastic) [1]. Extracellular hydrophobic barriers of lignin and suberin located in the endodermis are thought to restrict diffusion of water along the apoplastic path [2]. In the presence of such barriers, the cell to cell path is favored by water channels present in cell membranes called aquaporins [3].

Aquaporins (AQP) are present throughout the living kingdom at the exception of thermophilic Archaea and intracellular bacteria [4]. They belong to the Major Intrinsic Proteins (MIP) superfamily and are characterized by six transmembrane domains and two highly conserved Asn-Pro-Ala (NPA) motifs [5]. Other typical signatures of AQP consist in selectivity filters domains structuring the pore and composed of the aromatic/arginine (ar/R) motifs and the Froger’s residues [6,7]. In higher plants, AQP isoforms fall into five families comprising the Plasma membrane Intrinsic Proteins (PIP), the Tonoplast Intrinsic Proteins (TIP), the Nodulin26-like Intrinsic Proteins (NIP), the Small Intrinsic Proteins (SIP) and the uncharacterized (X) Intrinsic Proteins (XIP) [8]. Although plant AQP have been localized throughout the cell secretory system, PIP, NIP and XIP are preferential residents of the plasma membranes while TIP accumulate in the tonoplast and SIP in the endoplasmic reticulum [9–11]. Functional studies combined with modelling approaches demonstrated that, more than being permeable to water, AQP possess a wide range of selectivity profiles [12,13]. Some PIP isoforms are thereby permeable to H_2_O_2_ and CO_2_, some TIP isoforms to NH_3_ and urea, and some NIP to small organic solutes or mineral nutrients [14–18]. AQP possess wide range of physiological functions and have now been identified in a number of crop species such as rice, maize, tomato, cotton, sorghum, *Setaria italica*, watermelon or *Cannabis sativa* [19–24].

In plants, AQP are involved in water transport in both roots and shoots. In Arabidopsis for instance, isoforms AtPIP2-2 and AtPIP1-2 contributes to around 14% and 20% of the root osmotic conductivity and shoot hydraulic conductivity, respectively [25,26]. AQP also have important roles in plant growth, CO_2_ fixation, nutrient allocation, reproduction or biotic interactions [8]. In recent years, AQP functions in plant water relations have received more attention as a potential target for crop improvement, particularly to increase tolerance to drought [27–32]. For instance, AQP could regulate root water transport in order to match transpiration and therefore contribute to mitigate disparity between water supply and demand in soybean upon “atmospheric” water stress caused by high evaporative demand [33]. This assumption is supported by the increased AQP expression and root hydraulic conductivity upon transpiration demand in rice and grapevine [34,35]. Furthermore, overexpression of *SiTIP2-2* in tomato increased transpiration rate and was associated with improved fruit yield upon moderate soil water stress [36]. Conversely, in other crops more adapted to hot and dry climates such as pearl millet, lower transpiration under high vapor pressure deficit (VPD) has been proposed to be beneficial for tolerance to “atmospheric” water stress and potentially associated with AQP function in root hydraulics [30,37,38]. Therefore, AQP may be involved in different physiological mechanisms that determine the extent of water usage by the plant [39]. In fact, specific isoforms might be involved in different scenarios, calling for a better understanding of AQP family members and their functions in crops [28].

Pearl millet [*Pennisetum glaucum* (L) R Br.] is a key cereal for food security in arid and semi-arid regions [40]. However, its yield remain low and is often affected by climate unpredictability (heat waves and dry spells) that are forecast to worsen in future climate change scenarios [41,42]. The recent release of the pearl millet genome has opened new ways for functional genomic-based efforts aiming at improving its yield and tolerance to abiotic stresses [43]. In this study, we evaluated the potential links between AQP function in roots and water use, a drought tolerance-related trait, by measuring AQP contribution to Lpr and AQP genes expression in the roots of two pearl millet inbred lines contrasting for water use efficiency. To have more insights into the AQP isoforms contributing to root hydraulics, we characterized the entire AQP family in pearl millet using a genomic approach.

## Materials and methods

### Plant material and growth conditions

IP4952 and IP17150, two pearl millet inbred lines that are part of the Pearl Millet inbred Germplasm Association Panel (PMiGAP) were used in this study [43]. The water use efficiency of these two lines was characterized across two lysimeters experiments performed under well-irrigated conditions at the International Crops Research Institute for the Semi-Arid Tropics (ICRISAT, India) according to [44]. These experiments indicated that IP4952 and IP17150 displayed relatively low and high water use efficiency, respectively (2.35 versus 3.72) [45] These two lines were used for root hydraulic conductivity measurements and AQP expression analyses in which plant were grown in hydroponic conditions into a nethouse at the IRD/ISRA Bel Air research station (Dakar, Senegal; 14.701615 N – 17.425359 W). Plants were germinated in Petri dishes into a chamber at 37°C for two days in the dark. Plants were further exposed to light (37°C, 12h day/night cycle) for one day before being transplanted on top of a black mat covering a 30L container (45×40×24 cm containing 40 plants in total) filled with half strength Hoagland solution [46]. The system allowed roots to strictly develop in the nutrient solution without been exposed to light. Oxygen was provided to the roots through constant bubbling of the solution using an air pump.

### Root hydraulic conductivity and aquaporin contribution

Root hydraulic conductivity (Lpr) was measured in April-Mai 2019 (17/34°C and 34/100% in minimum/maximum temperature and humidity, respectively) from 9AM to 12PM on 15 days old plants grown in hydroponic conditions. The average temperature, relative humidity and VDP at the time of measurements were 24.4 ± 0.3°C, 75.1 ± 1.2% and 0.8 ± 0.1, respectively. Plants were grown sequentially to be analyzed at the same age in a randomized design taking into account the time of measurement. Lpr measurements were performed using a pressure bomb (model 1000, PMS instrument company, USA) according to [47] and [48]. Briefly, plants were inserted into the pressure chamber filled with nutrient solution complemented or not by 2mM of azide (NaN_3_). The hypocotyl was carefully threated through the silicone grommet of the pressure chamber lid while the intact root system was sealed into the chamber. Roots were pressurized with compressed air at 0.4 MPa for 5 min to equilibrate, followed by xylem sap collection at 0.1, 0.2 and 0.3 MPa for 5 min using pre-weighed 2 ml Eppendorf tubes filled with cotton placed on top of the stem. The mass of xylem sap exuded at each pressure was determined by weighing and used to calculate the xylem sap flow (slope of xylem sap weight at each pressure). After the measurements, roots systems were scanned to determine root surface area using WhinRhizo Pro version 2012b (Regent Instruments, Canada). Xylem sap flux was divided by root surface area to calculate Lpr. AQP contribution to Lpr was estimated using relative Lpr inhibition by azide calculated as:

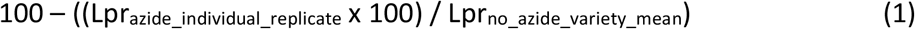

### Aquaporins genome-wide identification

A total of 772 AQP protein sequences from 19 plant species (*Arabidopsis thaliana, Beta vulgaris, Brachypodium distachyon, Cicer arietinum, Gossypium hirsutum, Glycine max, Hordeum vulgare, Linum usitatissimum, Musa acuminate, Panicum virgatum, Pennisetum glaucum, Populus tremula, Oryza sativa, Setaria italica, Solanum lycopersicum, Solanum tuberosum, Sorghum bicolor, Vitis vinifera* and *Zea mays*) were aligned against the pearl millet genome (ASM217483v2) and the non-assembled scaffolds [43] using tblastn with an e-value of 10^−5^ as initial cut-off to identify high scoring pairs. High scoring pairs were further filtered to keep those with a bit score ≥100. Hot-spots of high scoring pairs were identified and redundant high scoring pairs were filtered to keep those with highest bit-score for further analysis (S2 Table). The filtered high scoring pairs locations in the pearl millet genome were used to identify regions with homologies to AQP genes.

### Structural annotation

Correspondence between selected high scoring pairs and annotated genes in the pearl millet genome was determined. Potential AQP genes were identified and their genomic sequence ± 1000pb upstream and downstream of the start/end position was retrieved as well as the predicted gene structure [43]. When predicted AQP did not correspond to previously annotated genes, the GENSCAN Web Server (http://hollywood.mit.edu/GENSCAN.html) was used to predict the exon-intron structure of the genomic region. Putative AQP genomic sequences were aligned against the Plant EST (downloaded in August 2018) and the UniProt/Swiss-Prot plant protein (February 2016) databases and manually annotated using the Artemis software (version 17.0.1, Sanger Institute, UK; S3 Table). Annotation was confirmed by aligning reads from pearl millet transcriptomes [49] on the pearl millet genome using the Tablet software (version 1.19.09.03) [50] and coding and protein sequences were generated. AQP gene structure were further visualized using GSDS2.0 software [51].

### Sequencing

AQP genes sowing missing sequences in or bordering coding regions were resequenced (S3 Table) using genomic DNA or cDNA from Tift 23D2B1 (genotype used to draft the pearl millet whole genome sequence). DNA was prepared using DNeasy Plant mini extraction kit (Qiagen, Germany) while cDNA was prepared by first extracting RNA using the RNeasy Plant mini extraction kit (Qiagen, Germany) followed by DNAse treatment (RNase-free DNase set; Qiagen, Germany) and reverse transcription reaction (Omniscript RT kit; Qiagen, Germany) according to the manufacturer’s instructions. Corresponding DNA/cDNA fragments were amplified using the Phusion high-fidelity DNA polymerase (Thermo Scientific, USA), purified (Geneclean turbo kit, MP Biomedicals, USA) and sent for sequencing (Eurofins Genomics, Germany). Primers used for amplification are presented in S4 Table. Difficulties in amplifying the missing sequence of PgTIP5-1 were encountered. In that specific case, we used unpublished MINION reads (Mariac, Vigouroux, Berthouly-Salazar; unpublished) to complete its sequence. Missing nucleotides were added to the pearl millet genomic sequence and coding frame of the corresponding new protein was carefully checked.

### Identification of functional motifs and transmembrane domains

The NCBI conserved domain database (CDD) was used to identify NPA motifs and aromatic/arginine (ar/R) selectivity filters in the putative AQP protein sequences. Froger’s residues were identified according to [6]. The number and location of the transmembrane domains were studied using TMHMM (http://www.cbs.dtu.dk/services/TMHMM/), TMpred (https://embnet.vital-it.ch/software/TMPRED_form.html) and Phyre2 [52]. Protein sequences were aligned using the CLUSTALW alignment function in the Maga7 software (version 7.0.26) [53]. Alignments were colored using the Color Align Properties program (https://www.bioinformatics.org/sms2/color_align_prop.html). Conserved domains as well as transmembrane domains were further manually analyzed to detect sequence alterations. Three-dimensional geometry structure and pore morphology of PIP was obtained using the PoreWalker software (https://www.ebi.ac.uk/thornton-srv/software/PoreWalker/).

### Phylogenetic analysis

Phylogenetic analyses of *P. glaucum* AQP (PgAQP) was analyzed in relation to AQP identified in *A. thaliana* (AtAQP), *O. sativa* (OsAQP) and *P. tremula* (PtAQP) using the Mega7 software (version 7.0.26) [53]. PgAQP, AtAQP, OsAQP and PtAQP protein sequences were aligned using the CLUSTAW function and phylogenetic tree was built using the maximum likelihood method with 1000 reiterations. This allowed determination of the statistical stability of each node. Based on their position in the phylogenetic tree, PgAQP isoforms were classified into PIP, SIP, TIP and NIP families and named according to their close homologs.

### Expression profiling

*P. glaucum* PIP (PgPIP) genes expression were measured in root of 15 days old plants grown in hydroponic conditions using quantitative polymerase chain reaction (RT-PCR). Roots (seminal + one crown root) were sampled between 10AM to 12PM. Sampled roots were immediately frozen into liquid nitrogen and ground using a TissueLyser II (Qiagen). RNA and cDNA (from 1μg of RNA) were prepared using extraction kits as described above. RT-PCR was performed with 1μL of diluted cDNA (1:9) in a Brillant III ultra fast SYBRgreen QPCR master mix (Agilent Technologies, USA) using a StepOnePlus Real-Time PCR System (Applied biosystems, USA). Primers used to amplify the different PgPIP genes were checked for specificity and efficiency prior to the experiment (S5 Table). The pearl millet Ubiquitin gene (*Pgl_GLEAN_10001684*) was used as reference and PgPIP relative expression was calculated according to the delta-delta ct method.

PgAQP expression in shoots were retrieved from [49]. Data from leaves and inflorescence of ten open pollinated cultivated pearl millet varieties were used [49].

### Statistics

Statistical analyses were performed using R version 3.5.2 (R Development Core Team, 2018) using ANOVA (aov script) to detect significant differences. Least Significant Difference (LSD) test within the Agricolae package was used to group differences in letter classes.

## Results

### Pearl millet aquaporin contribution to root hydraulic conductivity

In order to evaluate if AQP function in root radial water flow could be associated with water use in pearl millet, we measured root hydraulic conductivity (Lpr) in IP4952 and IP17150, previously described as low and high water use efficiency lines respectively. IP4952 showed 1.4 times higher Lpr than IP17150 (1.30E-07 ± 2.36E-08 versus 9.27E-08 ± 8.33E-09 m^3^ m^−2^ s^−1^ MPa^−1^ respectively; n=15) but this difference was not significant (*p* = 0.148; Fig 1 and S1 Table). In both IP4952 and IP17150, treatment with azide, an inhibitor of AQP activity, led to significant Lpr reduction to 2.00E-08 ± 2.44E-09 and 2.19E-08 ± 2.32E-09 m^3^ m^−2^ s^−1^ MPa^−1^ respectively (*p* < 0.001; S1 Table). This effect of azide application on Lpr was mostly reversible after treating the same roots with azide-free solution (S1 Fig). AQP contribution to Lpr was significantly higher in IP4952 (low water use efficiency) as compared to IP17150 (high water use efficiency; 84.64 ± 1.98% versus 76.40 ± 2.61%, *p* < 0.05; S1 Table). These data indicate that AQP could contribute more than 75% to Lpr in pearl millet.

**Fig 1.**
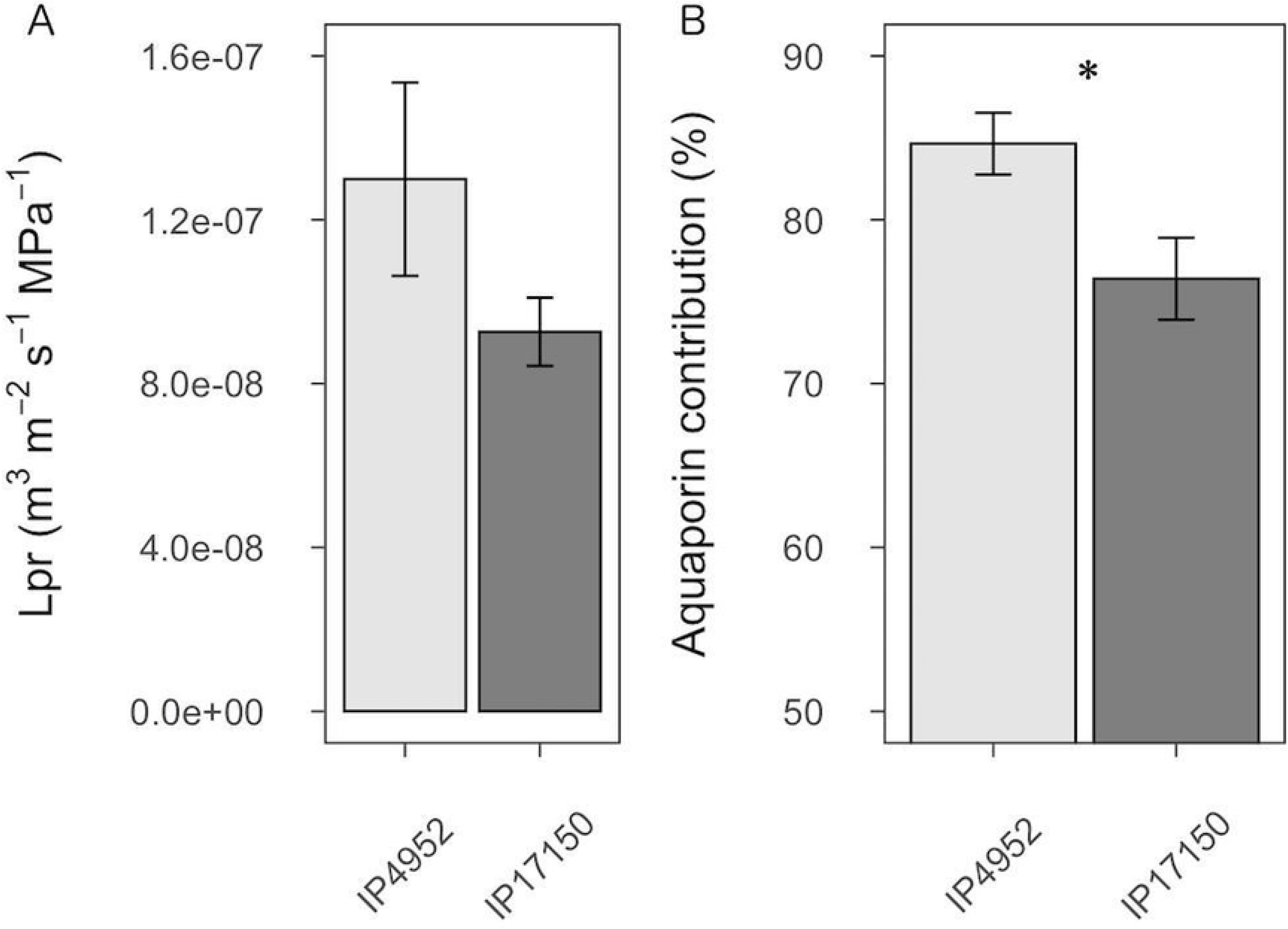
Root hydraulic conductivity and aquaporin contribution in roots of IP4952 and IP17150. Root hydraulic conductivity (Lpr) were measured in plants grown in hydroponic conditions between 9AM to 12PM in presence or not of 2mM azide. (A) Lpr values measured in absence of azide. (B) Aquaporin contribution to Lpr was calculated as relative Lpr inhibition by azide. Bars represent mean values ± se of n=10-15 plants. *: *p* < 0.05.

### Aquaporin identification and annotation

To have more insight in the AQP isoforms contributing to Lpr in pearl millet, we characterized the AQP genes family in the pearl millet genome. We blasted 772 AQP from the PIP, TIP, SIP, NIP and XIP families identified in 19 different species on the pearl millet reference genome (chromosome assembly and scaffolds) [43]. A total of 7005 sequences with bit score >100, representing 50 specific hits spread on the genome were identified (S2 Table). Forty-seven of the hits fall into previously annotated genes, one fall in a non-annotated part of the genome on chromosome 5, and two in non-assembled parts of the genome (scaffold763 and scaffold8428).

Manual *de novo* annotation of the 50 putative AQP genes allowed the identification of eight genes with no start or with early stop codon in the first exon that were classified as pseudo-genes (S3 Table). Nine genes did not encode AQP isoforms but zinc-finger protein/LRR receptor-like serine-threonine protein kinase families or DEAD-like helicase-N superfamily, respectively. The absence of AQP signature domains (NPA and Ar/R motifs) in their protein sequence confirmed their non-affiliation to the AQP family. A number of genes showed an excessive number of exons or longer first exon and were re-annotated on the basis of alignment with transcriptome sequences or with close protein homologs (Uniprot/Swiss-Prot blast results) and presence of AQP isoform conserved domains. In addition, ten genes showed missing sequences in coding regions that were completed after sequencing. Overall, 33 *Pennisetum glaucum* AQP (PgAQP) genes were identified in the pearl millet genome, among which sixteen were *de novo* annotated (Table 1).

**Table 1.**
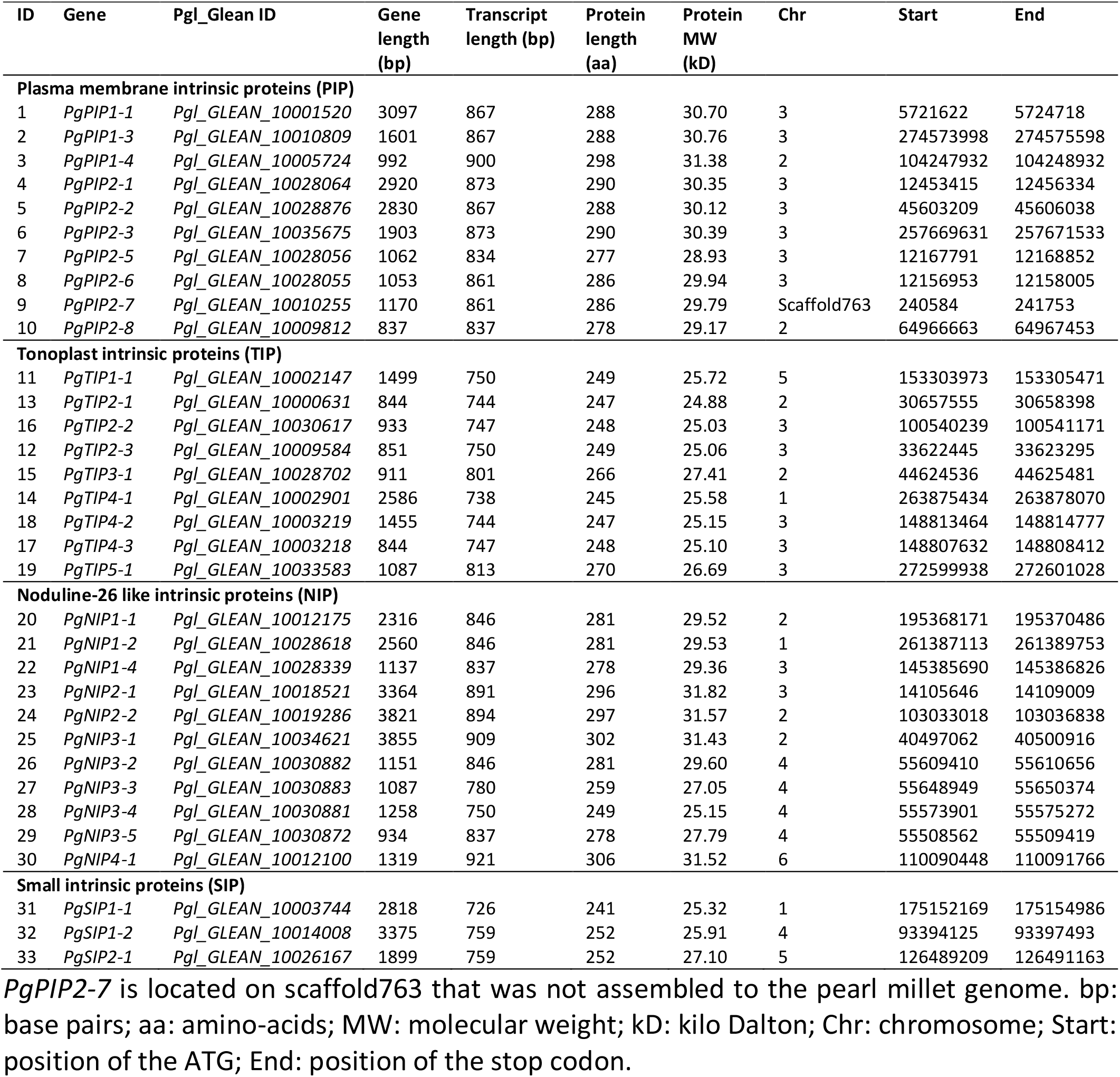
Description and distribution of the aquaporins genes identified in pearl millet.

### Pearl millet aquaporins phylogenic analysis

To classify the PgAQP into families and name them, a phylogenetic tree was built using the PgAQP protein sequences along with protein sequences from *Arabidopsis thaliana*, *Oryza sativa* and *Populus tremula* (Fig 2). The PgAQP were named according to their grouping into families (PIP, TIP, SIP or NIP) and close homologs (Table 1). Ten isoforms showed homologies to the PIP family with three isoforms falling in the PIP1 sub-family (PgPIP1-1, PgPIP1-3 and PgPIP1-4) and seven isoforms falling in the PIP2 sub-family (PgPIP2-1, PgPIP2-2, PgPIP2-3, PgPIP2-5, PgPIP2-6, PgPIP2-7 and PIP2-8). Nine isoforms from the *P. glaucum* TIP family (PgTIP), eleven isoforms from the *P. glaucum* NIP family (PgNIP) and three isoforms from the *P. glaucum* SIP family (PgSIP) were further identified. No isoforms from the XIP family were identified in pearl millet. This analysis further confirmed the classification of PgPIP1-1, PgPIP2-1, PgPIP2-3, PgPIP2-6, PgTIP1-1 and PgTIP2-2 cloned by [30]. However, PgPIP1-2 from [30] was renamed as PgPIP1-3 in this study.

**Fig 2.**
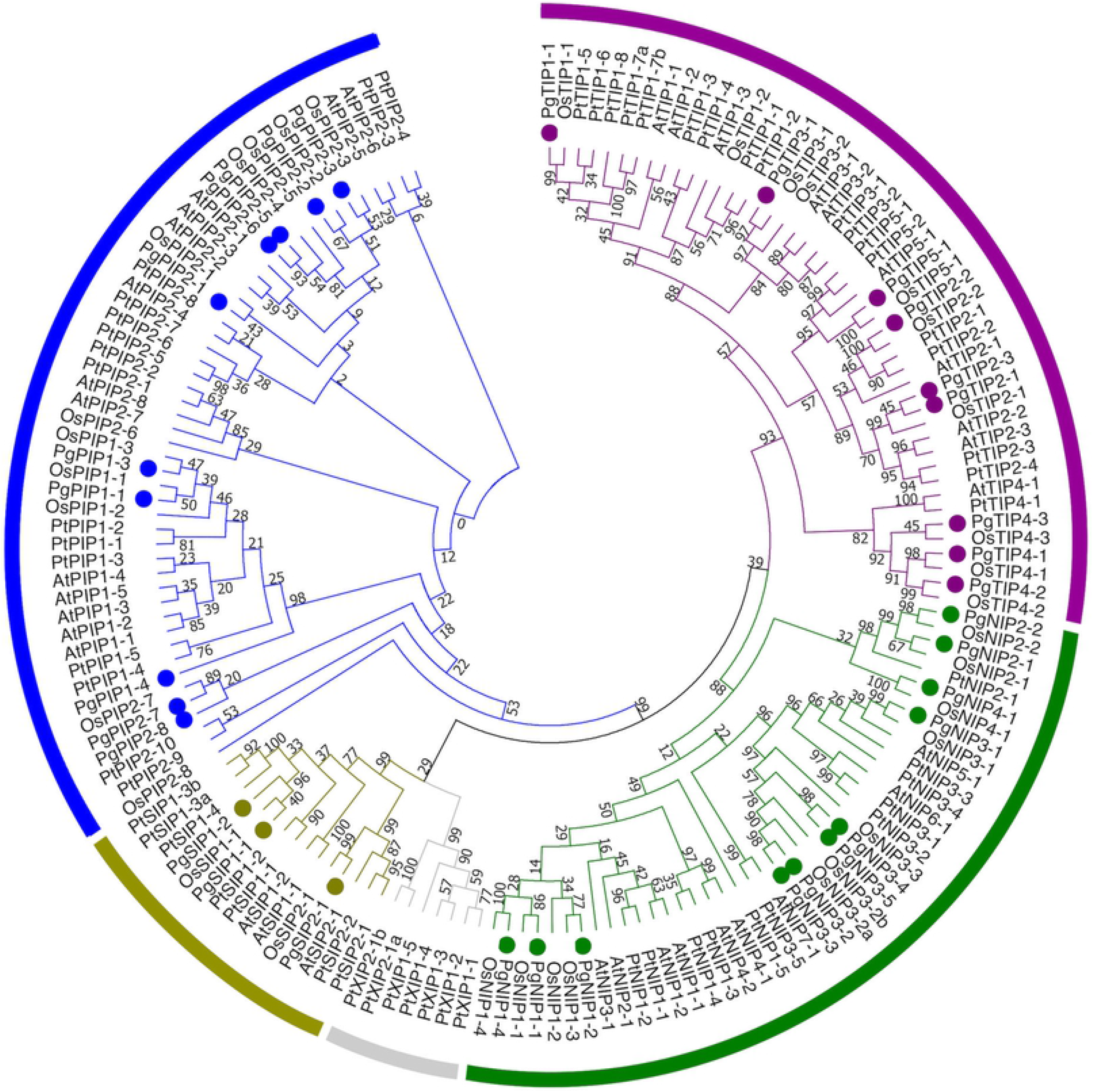
Phylogenetic relationship among aquaporins isoforms from pearl millet, Arabidopsis, rice and poplar. Tree was generated using the Maximum Likelihood method with 1000 reiterations in MEGA7. Bootstrap values above 50% are represented. The PIP, TIP, SIP, NIP and XIP family clades are represented by blue, grey, purple, green and red, respectively. Pearl millet sequences are indicated by colored dots.

PgAQP genes distribution in the pearl millet genome denoted that most of them were localized on chromosome 3 while none were localized on chromosome 7 (Fig 3). Two hot-spots of PgAQP genes were observed, one located in a region of 11899bp on chromosome 3 containing *PgPIP2-1*, *PgPIP2-5* and *PgPIP2-6* and the other in a region of 141812bp on chromosome 4 containing *PgNIP3-2*, *PgNIP3-3*, *PgNIP3-4* and *PgNIP3-5*.

**Fig 3.**
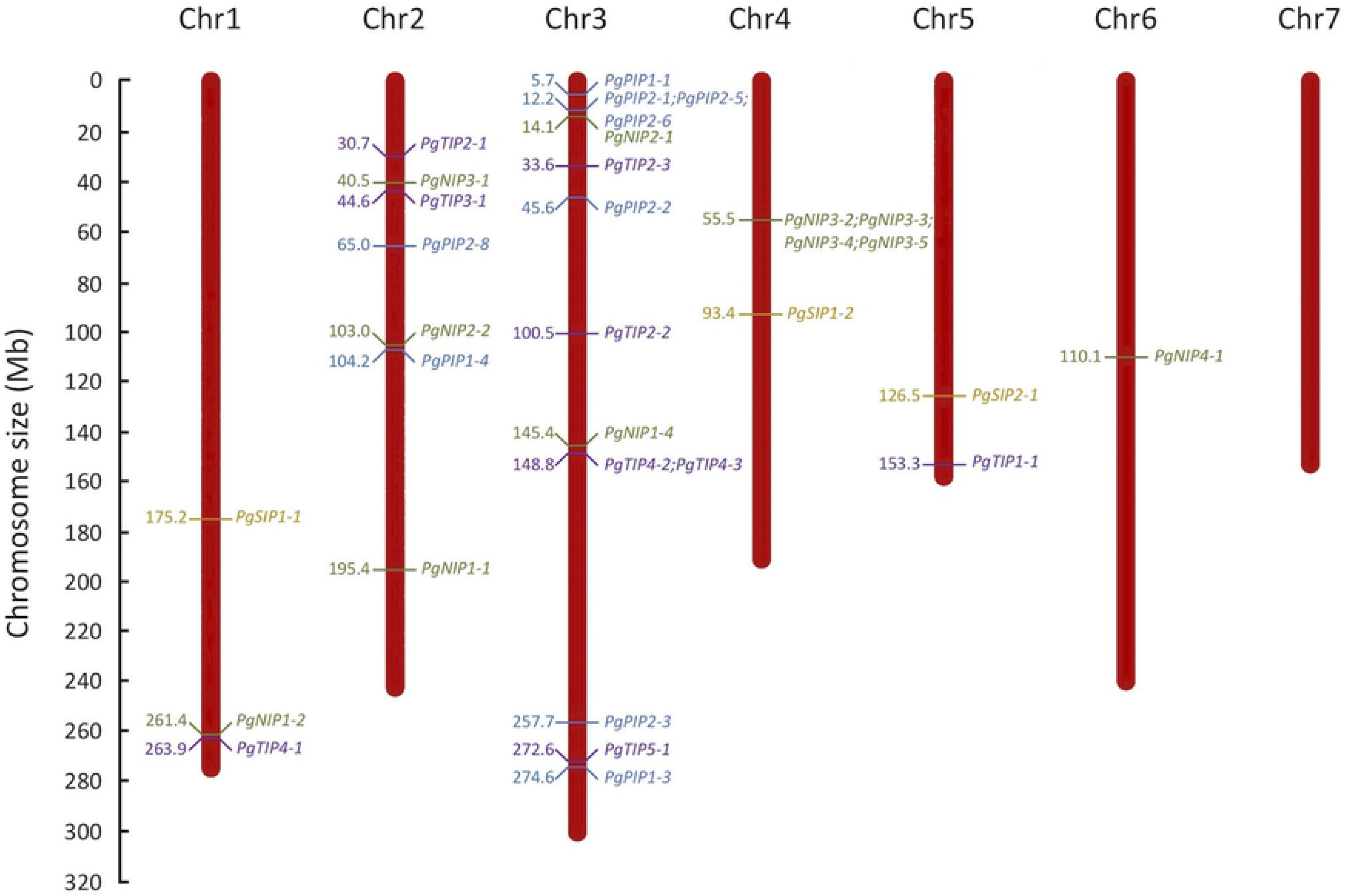
Distribution of aquaporin genes in the pearl millet genome. The seven chromosome (Chr) from the pearl millet genome are represented according to their size in megabase (Mb). Positions of PIP, TIP, SIP and NIP genes are represented in blue, purple, orange and green, respectively. *PgPIP2-7* which is located on scaffold763 is not represented.

### Aquaporin gene structure in pearl millet

PgAQP genes showed large variation in gene length (ranging from 837bp for *PgPIP2-8* to 3855bp for *PgNIP3;1*; Table 1). Transcript length were less variable and relatively conserved within families with lengths of around 800-900bp for the PgPIP and PgNIP genes, 750-800bp for the PgTIP genes and 700-750bp for the PgSIP genes.

To investigate associations between phylogenetic classification and gene structure, the intron-exon organization of the PgAQP genes was analyzed (S2 Fig). PgAQP genes displayed between one (*PgPIP2-8*) to five exons (*PgNIP2-1*, *PgNIP2-2* and *PgNIP4-1*). Except for the NIP family, intron-exon organization was generally conserved within families with 5/9 PgPIP genes displaying 3 exons, 6/9 PgTIP genes displaying 2 exons and all PgSIP genes displaying 3 exons, supporting their phylogenetic distribution. *PgPIP2-5/PgPIP2-6* and *PgNIP3-2/PgNIP3-3* that were found to be close homologs in the phylogenetic analysis (Fig 2) and closely located in the pearl millet genome (Fig 3) showed similar gene and transcript length (Table 1) as well as gene structure (S2 Fig). PgAQP coding regions encoded proteins with length varying between 250 to 300 amino acids, with molecular weight of around 30kD for the PgPIP and PgNIP and 25kD for the PgTIP and PgSIP isoforms (Table 1).

### Pearl millet AQP conserved domains

To investigate polymorphisms in the PgAQP isoforms conserved motifs that could underlay potential changes in structural and substrate selectivity, analyses of conserved and transmembrane domains were performed. The conserved domain analysis showed that all PgAQP isoforms belonged to the MIP super-family and displayed typical double NPA motifs (S6 Table and Table 2). Although some polymorphisms in the NPA motifs were observed in some isoforms (particularly from the NIP family) the subsequent amino-acids were of the same chemical properties (generally neutral and non-polar; Table 2, Fig 4 and S3-5 Fig). Ar/R selectivity filters and Froger’s residues known as AQP markers were well conserved in the PgPIP isoforms except for the Froger’s residue on position 1 (P1; Table 2, Fig 4). More polymorphisms were observed in these residues for the PgTIP, PgNIP and PgSIP isoforms although the ar/R residue on Loop E (R on LE2) and the Froger’s residues at positions 3 (A) and 4 (F/Y) were well conserved across all isoforms (Table 2).

**Table 2.**
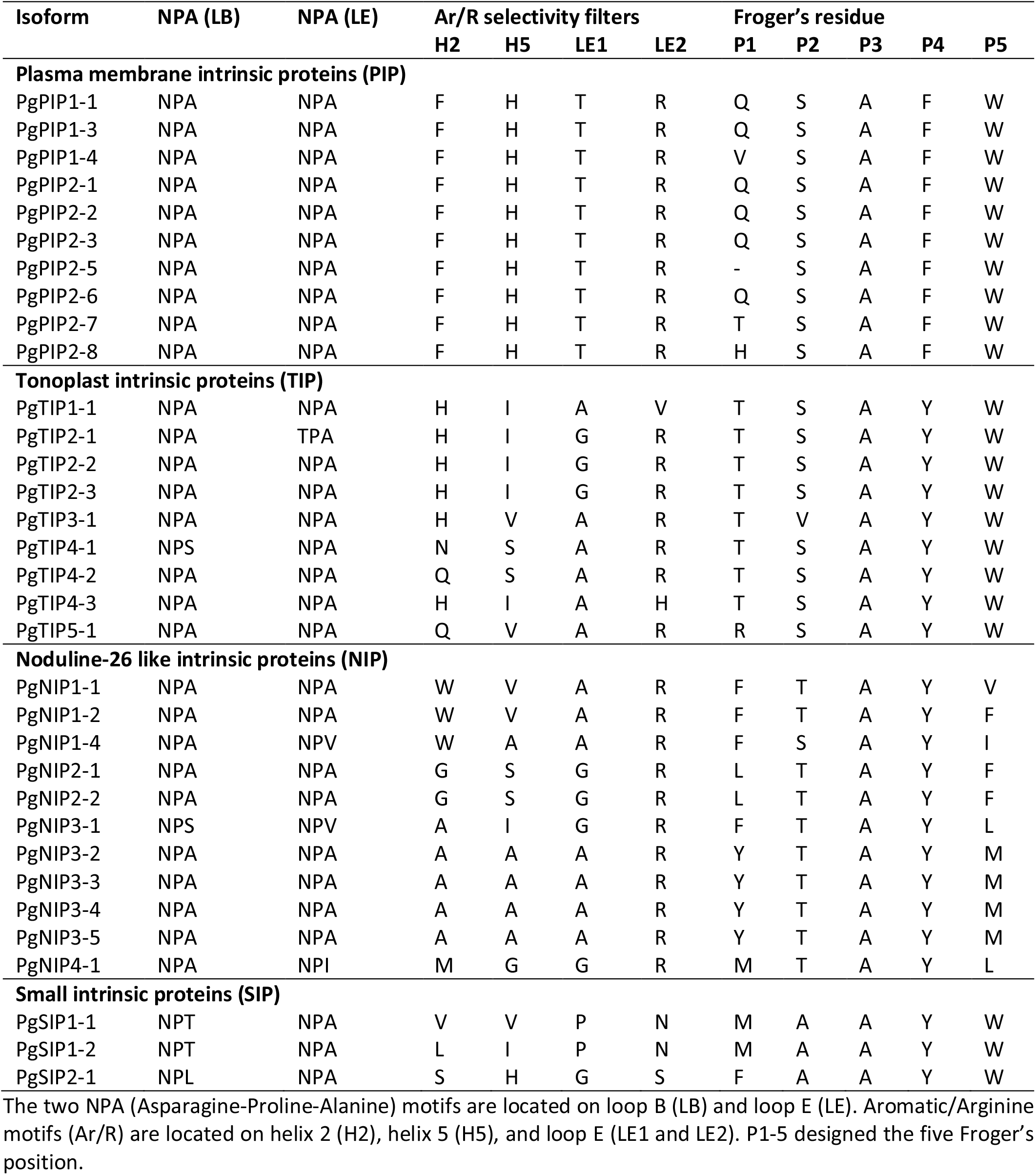
Amino-acids residues in the conserved domains of the pearl millet aquaporins isoforms.

**Fig 4.**
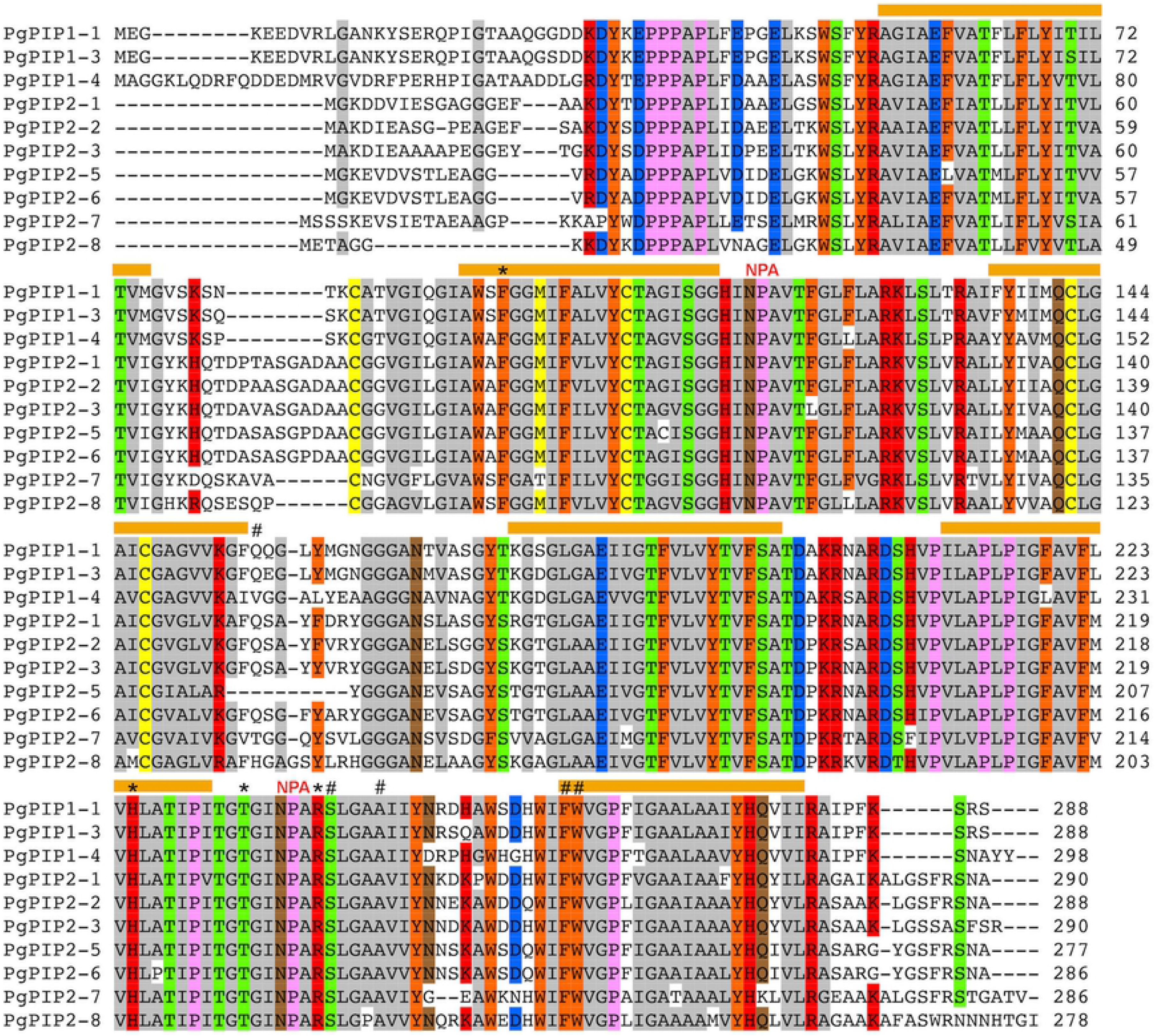
Conserved domains and membrane topology of the PIP isoforms from pearl millet. Alignment of the PIP isoforms were obtained using ClustalW in Mega7. Sequence identities and similarities (80%) are highlighted in colors. The transmembrane domains are represented by orange bars with the N-terminal and C-terminal ends of the protein located in the cytosol. NPA: Asparagine-Proline-Alanine motifs; *: Aromatic/Arginine selectivity filters. #: Froger’s residues.

Transmembrane domains analyses using three different prediction softwares suggested a high probability for all identified PgAQP to display six transmembrane domains with cytoplasmic N-terminal and C-terminal ends as typically observed for AQP (S7 Table). Predictions of the 3-dimensional geometry structure and pore morphology suggested a continuous pore that runs longitudinally across both sides of the membranes for all PgPIP (S6 Fig). Two deviations in the pore center were typically observed at both extremities illustrating the two constraints caused by the NPA motifs structured as alfa-helixes and acting as selectivity filters.

### Aquaporin expression profiling in pearl millet

Because of their localization at the plasma membrane, PIP isoforms play major roles in root hydraulic conductivity. To infer PgPIP isoforms putative importance to Lpr in pearl millet, we analyzed PgPIP gene expression pattern in roots of IP4952 and IP17150 using quantitative PCR. PgPIP genes were generally more expressed in IP4952 as compared to IP17150 (Fig 5). In both lines, the most expressed genes in roots were *PgPIP1-1*, *PgPIP1-3*, *PgPIP1-4*, and *PgPIP2-3* while *PgPIP2-5*, *PgPIP2-6*, *PgPIP2-7* and *PgPIP2-8* were lowly expressed. *PgPIP1-3* and *PgPIP1-4* were the only genes significantly differentially expressed between both lines.

**Fig 5.**
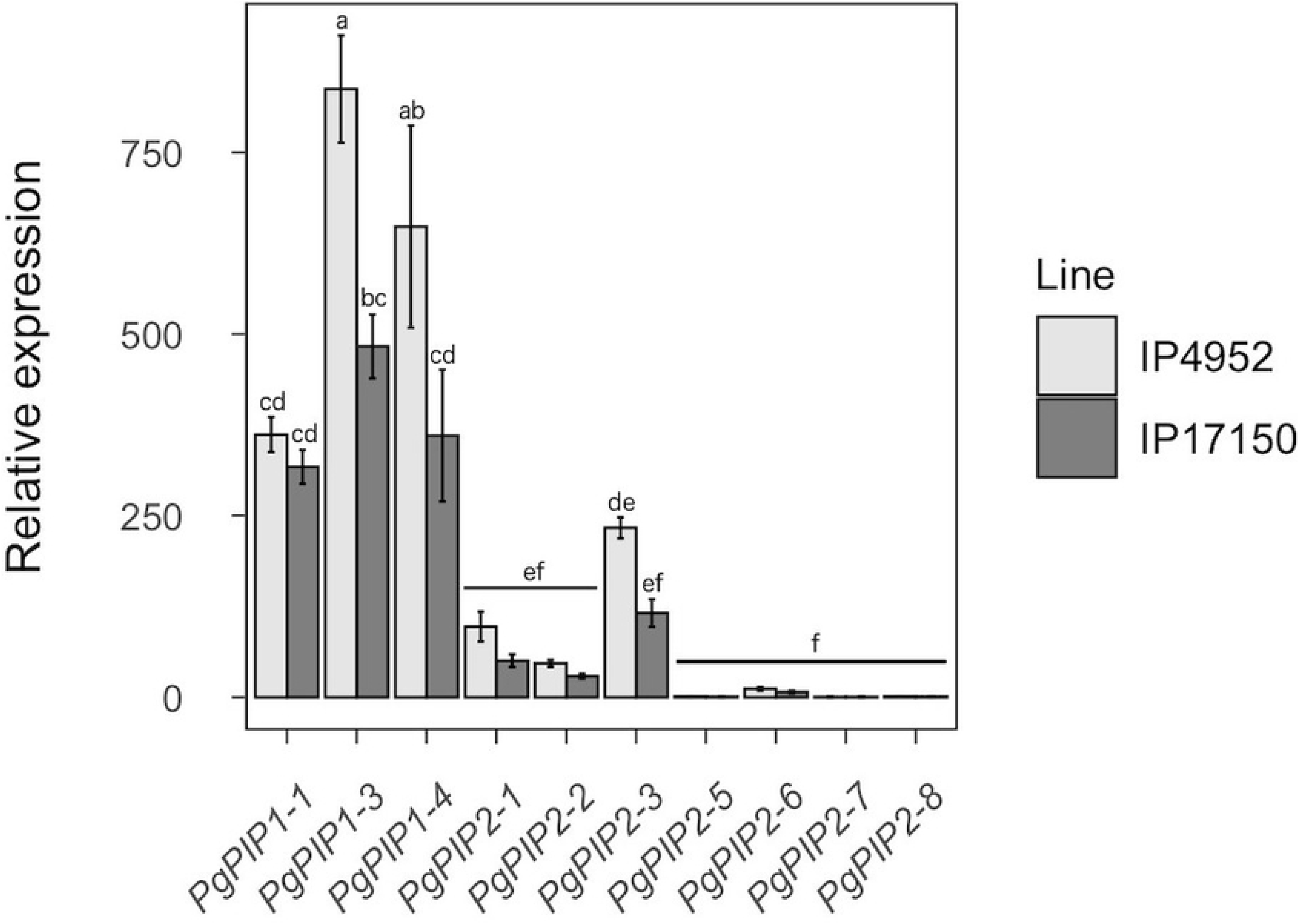
Relative expression of PIP genes in roots of IP4952 and IP17150. Transcript abundance of each PIP genes were measured between 9AM and 12PM on plants grown in hydroponic conditions and normalized to the expression of *PgPIP2-5* in IP4952. Bars show mean values ± se of n=6-8 biological replicates, each with technical triplicates. Letters indicate different significance groups.

Expression of PgPIP in shoots (leaves and inflorescence) were retrieved from [49]. Transcriptomic analyses from ten pearl millet varieties suggest that *PgPIP1-4* and *PgPIP2-3*, two of the most expressed genes in roots, are lowly expressed in shoots (S7 Fig). Conversely, *PgPIP1-1* and *PgPIP1-3* are highly expressed in roots and shoots.

## Discussion

Here, we studied the role of AQP in root water transport in pearl millet, a heat and drought-adapted crop. Root hydraulic conductivity (Lpr) varied around 1E-07 m^3^ m^−2^ s^−1^ MPa^−1^, in a range that has been previously reported in other plants [54]. Root treatment with a common AQP inhibitor (azide) suggested that AQP contribute up to 84% to the Lpr in pearl millet (S1 Table). This figure is higher than what has been observed in Arabidopsis (57 to 64%) [55] and rice (42 to 79%) [56]. However, as complete reversion of Lpr inhibition by azide could be observed (S1 Fig), we do not think AQP contribution might have been over-estimated due to azide secondary effects. Hence, our results suggest that PgAQP are major regulators of root water flow in pearl millet.

We used the pearl millet genome sequence to characterize the AQP gene family and identified thirty-three putatively functional AQP isoforms (based on conserved domains and protein topology) that belonged to the PIP (10), TIP (9), SIP (3) and NIP (11) families. No XIP were identified in pearl millet which confirm the absence of isoforms from this family in the monocotyledon clade [8]. The number of identified AQP in pearl millet is similar to what has been observed in *A. thaliana* (35) [57], rice (33) [22] and maize (31) [19]. Interestingly, AQP genes were over-represented on Chromosome 3 with fourteen genes (Fig 3). *PgPIP2-5* and *PgPIP2-6* closely located in Chromosome 3 are phylogenetically related and possess similar gene structure (Fig 3 and S2 Fig) suggesting possible tandem duplication events in this region [58]. Similarly, *PgNIP3-2* and *PgNIP3-3* located on Chromosome 4 may be the result of duplication events. Furthermore, *PgPIP2-7* was identified in scaffold763 suggesting that genes may be missing on parts of the pearl millet genome assembly.

The selectivity of plant AQP is defined by amino-acids structuring the pore that constitute their signatures. Among these amino-acids, the NPA motifs on loops B and E contribute with the dipole moment of the α-helices to prevent proton permeation [59,60]. These motifs were strictly conserved in the PgPIP but showed some polymorphisms for other isoforms (Table 2). However, these substitutions did not drastically change the positive electrostatic potential at the NPA motifs suggesting that proton exclusion from AQP pore is conserved in pearl millet. Furthermore, the Ile preceding the Froger’s residue P4 and P5 at the end of Loop E, shown to be essential for CO_2_ transport in PIP [61], is conserved in the PgPIP (Fig 4).

The ar/R motifs, composed of four amino-acids forming a constriction at the extracytosolic entry of the pore, represent the main selectivity filter. Modelling approaches based on ar/R signatures were used to predict permeability of plant AQP [12,13]. For instance, the F-H-T-R signature observed in the PgPIP (Table 2) which seems strictly conserved across PIP from different species has been associated with water and H_2_O_2_ permeability [14,25,26,62]. The H-I-G-R signature observed in PgTIP2-1, PgTIP2-2 and PgTIP2-3 that is conserved in TIP2 from Arabidopsis, maize and rice supposedly allow permeability to water, NH_3_, urea and H_2_O_2_ while the H-I-A-V signature observed in the unique PgTIP1 (PgTIP1-1) may allow NH_3,_ urea and H_2_O_2_ but restrict water permeation [63–65]. In PgNIP, the W-V-A-R signature have been associated with water, NH_3_ and H_2_O_2_ transports while the A-I/A-G/A-R signatures were restricted to water and NH_3_. Interestingly, PgNIP2-1 and PgNIP2-2 showed similar ar/R signature (G-S-G-R) than OsNIP2-1 that is permeable to silicon [18] and possess precise spacing of 108 amino-acids between the two NPA motifs supposed as essential for silicon permeability [66].

AQP in plants are expressed from root to leaf tissues, including inflorescence and pollen. Some PgPIP genes show tissue-specific expression with *PgPIP1-4* and *PgPIP2-3* being more specifically expressed in roots while *PgPIP2-1* is more specifically expressed in shoots and *PgPIP1-1* and *PgPIP1-3* are expressed in both tissues (Fig 5 and S7 Fig). It has been shown that PIP isoforms agglomerate as tetramers in the plasma membrane, each monomer forming functional units. Functional studies as well as protein-protein interactions studies suggest that PIP tetramers can be formed of heteromers of PIP1 and PIP2 with distinct functional properties depending on the isoform combination [67–70]. Based on PgPIP gene expression in pearl millet, PgPIP1-1, PgPIP1-3 and PgPIP1-4 might interact with PgPIP2-3 to form heteromers in roots while PgPIP1-1, PgPIP1-3 and PgPIP2-1 might form different combinations of heteromers in shoots.

Intraspecific diversity in AQP isoform expression have been observed in rice and Arabidopsis [55,56,71,72]. In our study, diversity in expression pattern of PgAQP genes were observed between two pearl millet inbred lines with different water use strategies (Fig 5). IP4952 that showed significantly higher AQP contribution to Lpr as compared to IP17150 also showed significantly higher *PgPIP1-3* and *PgPIP1-4* gene expression. These results suggest that differences in expression of these AQP genes can reflect differences in AQP contribution to Lpr in pearl millet. These observations are in line with results from [34] showing that the expression of *VvPIP1-1* is associated to root hydraulics and response to water stress in two isohydric and anisohydric grapevine (*Vitis vinifera*) cultivars. Transpiration response to high VPD in four pearl millet recombinant inbred lines from a high resolution cross have been linked to PgPIP gene expression in roots [30]. These authors suggested that a down-regulation of PgPIP genes under high VPD induced reduction in transpiration and water savings. Our results showing reduced expression of *PgPIP1-3* and *PgPIP1-4* and AQP contribution to Lpr in IP17150, the line with higher water use efficiency, support the observations of [30]. Overall, expression profiling suggests that APQ may have different physiological functions across the pearl millet plant and contribute to its response to the environment. However, expression alone is certainly not fully representative of AQP function due to the many post-translational regulations affecting their activity [8]. Further investigations are needed to better understand the links between reduction in transpiration under high VPD, improved water use efficiency and AQP function in roots.

Pearl millet is a drought-adapted crop that will play a major role in the adaptation of agriculture to future climate in arid and semi-arid regions of Africa and India. Here, we provide a comprehensive view of the AQP genes and isoforms present in pearl millet as well as their contribution in root radial water transport. We confirmed the presence of selectivity filters suggesting permeability to water in the PgAQP and point isoforms PgPIP1-3 and PgPIP1-4 as potential main contributors of root water transport in pearl millet. The function of these isoforms may be subjected to natural diversity and associated with plant water use strategies as suggested by their differential expression in pearl millet lines contrasting for Lpr and water use efficiency. Therefore, our study supports a potential role for AQP in regulating pearl millet hydraulics and potentially adaptation to challenging environmental conditions.

## Acknowledgments

The authors are grateful to Dr Prakash Gangashetty (ICRISAT, Niger) for providing seeds of the inbred lines used in this study and Dr Kwanho Jeong (IRD, France) for his kind support in the preparation of this manuscript. The authors acknowledge the IRD iTrop HPC (South Green Platform) at IRD Montpellier for providing HPC resources that have contributed to the research results reported in this paper (https://bioinfo.ird.fr/- http://www.southgreen.fr).

## Author contributions

AG, CTD and LL designed the study. AG, PA, CTD, CFC and CM performed the experiments. AG, CTD, PA, CFC, CM, PG, VV, YV and LL analyzed the data and discussed the results. AG and LL wrote the paper. All authors read and approved the manuscript.

## Supporting information

**S1 Table. Root hydraulic conductivity (Lpr) and aquaporin (AQP) contribution in IP4952 and IP17150.**

**S2 Table. High scoring pairs with highest bit score at the fifty hot-spots.**

**S3 Table. Functional annotation of the pearl millet genomic regions corresponding to hotspots of High Scoring Pairs.**

**S4 Table. Primers used for genomic DNA (gDNA) or complementary DNA (cDNA) amplification of aquaporins showing missing sequence.**

**S5 Table. Primers used for quantitative RT-PCR.**

**S6 Table. Analysis of aquaporin conserved domains in pearl millet.**

**S7 Table. Transmembrane domain analysis of aquaporins in pearl millet.**

**S1 Fig. Reversion of azide-induced root hydraulic conductivity inhibition.**

**S2 Fig. Structure of pearl millet aquaporins genes.**

**S3 Fig. Conserved domains and membrane topology of the TIP isoforms from pearl millet.**

**S4 Fig. Conserved domains and membrane topology of the NIP isoforms from pearl millet.**

**S5 Fig. Conserved domains and membrane topology of the SIP isoforms from pearl millet.**

**S6 Fig. Cavity features of PgPIP isoforms.**

**S7 Fig. Expression pattern of PgPIP isoforms in shoots of pearl millet.**

